# Transcranial electrical stimulation over premotor cortex mimics attentional modulation of visual processing

**DOI:** 10.1101/2023.08.01.551431

**Authors:** Jonas Misselhorn, Marina Fiene, Jan-Ole Radecke, Andreas K. Engel, Till R. Schneider

**Affiliations:** Department of Neurophysiology and Pathophysiology, University Medical Center Hamburg-Eppendorf, 20246 Hamburg, Germany; Department of Psychiatry and Psychotherapy, University of Lübeck, 23562 Lübeck, Germany; Center of Brain, Behavior and Metabolism (CBBM), University of Lübeck, 23562 Lübeck, Germany

## Abstract

Attentional control over sensory processing has been linked to neural alpha oscillations and related pulsed inhibition of the human cortex. Despite the wide consensus on the functional relevance of alpha oscillations for attention, precise neural mechanisms of how alpha oscillations shape perception and how this top-down modulation is implemented in cortical networks remain unclear. Here, we tested the hypothesis that alpha oscillations in premotor cortex are causally involved in top-down regulation of visual cortex responsivity to contrast. We applied intermittent transcranial alternating current stimulation (tACS) over bilateral premotor cortex to manipulate attentional preparation in a visual discrimination task. tACS was applied at 10 Hz (alpha) and controlled with 40 Hz (gamma) and sham stimulation. Importantly, we used a novel linear mixed modeling approach for statistical control of neurosensory side-effects of the electric stimulation. We found a frequency-specific effect of alpha tACS on the slope parameter, leading to enhanced low-contrast perception and decreased perception of high-contrast stimuli. Side-effects affected both threshold and slope parameters, leading to high variability in parameter estimates. Controlling the impact of side-effects on psychometric parameters by linear mixed model analysis reduced variability and clarified the existing effect. We conclude that alpha tACS over premotor cortex mimicked a state of increased endogenous attention potentially by modulation of fronto-occipital connectivity in the alpha band. We speculate that this network modulation allowed for improved sensory readout from visual cortex which led to a decrease in psychometric slope, effectively broadening the dynamic range for contrast perception.

**Significance statement:** Attention is fundamental to voluntary control of perception and behavior. Yet, despite extensive scientific efforts, precise underlying neural mechanisms remain elusive. We contribute to this ongoing discussion by providing evidence for a vital role of frontal alpha oscillations in regulating the responsivity of visual cortex. By controlled neuromodulation with intermittent transcranial alternating current stimulation (tACS), we show that alpha tACS modulates psychometric properties of visual contrast perception. This study fills an important gap between work on alpha oscillations in spatial attention and studies on the psychometrics of attention. Furthermore, we pioneered an approach for the statistical control of tACS side-effects with linear mixed modeling and thereby add to the ongoing debate on outcome variability in studies using transcranial neurostimulation methods.

## Introduction

Theories of attention have evolved around the idea that sensory input can be voluntarily enhanced or attenuated in order to filter behaviorally relevant information (Broadbent, 1958; Treisman, 1964; Desimone, 1998). On a neurophysiological level, attention has been reliably related to alpha oscillations (Klimesch et al., 2007; Jensen and Mazaheri, 2010; Foxe and Snyder, 2011; Bonnefond et al., 2017). Specifically, alpha power modulations have been proposed to implement a gating mechanism that might route cortical information flow according to attentional demands (Jensen and Mazaheri, 2010). For example, spatial attention has been repeatedly shown to rely on differential amplitude enhancement or dampening of alpha oscillations, with stronger alpha activity in cortical regions receiving input from unattended spatial locations (van Dijk et al., 2008; Ikkai et al., 2016; Keefe and Störmer, 2021; Wöstmann et al., 2021; Radecke et al., 2023). Similarly to space, alpha amplitudes differ between sensory streams if one modality is attended at the expanse of the others (Foxe et al., 1998; Mazaheri et al., 2014). It was concluded that alpha oscillations likely reflect a general mechanism of pulsed inhibition (Jensen and Mazaheri, 2010) capable of shaping sensory processing by both local inhibition (Haegens et al., 2011) and cortical routing of information flow (van Dijk et al., 2008; Zhigalov and Jensen, 2020).

In a complementary approach, formal modeling of the effects of attention led to the proposal that neural response normalization might represent an underlying canonical neural computation (Reynolds and Heeger, 2009). Accordingly, neural population responses result from an initial afferent bottom-up drive that is scaled by top-down attentional modulation and normalized by a divisive input representing lateral inhibition in neural micro-circuits (Carandini and Heeger, 2012). A crucial advantage of using these models for human research is that, by measuring low-level psychophysics (e.g., slope and threshold for contrast perception), behavioral measures can be used to infer the underlying neural computation (Pestilli et al., 2009). A linking observation between formal modeling and the neurophysiology is that the alpha oscillation amplitude is held to shorten the “duty cycle” of gamma activity, and thereby to increase sensory thresholds (Jensen and Mazaheri, 2010). The psychometric threshold for visual detection has indeed been shown to be decreased by covert attention (Cameron et al., 2002). Also, it has been shown that the amplitude of alpha oscillations can be linked to the slope of the psychometric function (Chaumon and Busch, 2014). In this study, the amplitude of pre-stimulus alpha activity was indicative of the slope of the measured psychometric function. The authors interpreted this finding as evidence for fluctuations of response gain, such that the input/output relation of visual cortex was adapted in response to top-down influence. A likely candidate region providing these top-down signals are the human frontal eye fields located in premotor cortex that have been shown to play a key role in endogenous, but not exogeneous attention (Fernández and Carrasco, 2020; Fernández et al., 2023; Radecke et al., 2023). Furthermore, it was demonstrated that alpha oscillations in premotor cortex are involved in the top-down control of attention regulating perceptual gain (Misselhorn et al., 2019a).

Here, we set out to causally test the idea that preparatory top-down influence on visual processing rooted in premotor cortex (1) communicates through alpha oscillations, and (2), that this influence on visual processing is reflected as a change in psychometric slope. To this end, we applied intermittent transcranial alternating current stimulation (tACS) over the bilateral premotor cortex while measuring psychometric functions for visual contrast perception. Stimulation at alpha frequency (10 Hz) was controlled with gamma (40 Hz) and sham stimulation. In order to account for potential transcutaneous and transretinal side effects of tACS that might confound perception and behavior (Schutter, 2016; Asamoah et al., 2019), we introduce a novel linear mixed model analysis approach. We hypothesized that intermittent tACS would interfere with attentional preparation of visual perception and that only alpha stimulation would affect the slope of the psychometric function.

## Methods

### Participants

We enrolled 18 healthy young university students into the study (mean age 24.83 ± 3.75, 12 female, 6 male). All participants had normal or corrected-to-normal vision and reported no history of neuro-psychiatric disorders. All participants received monetary compensation for participation in the study and gave written informed consent in line with the Declaration of Helsinki prior to participation. The study was approved by the local ethics committee of the Medical Association of Hamburg.

### Experimental procedure

Participants were seated in a sound-attenuated and electrically shielded chamber. While preparing tACS electrodes mounted in an EEG cap (Easycap, Germany), participants were acquainted with the visual task. Thereafter, three blocks of identical psychometric testing were recorded under tACS application (sham, alpha, gamma; counterbalanced across subjects). We used an intermittent electrical stimulation protocol to restrict tACS to the period of attentional preparation and before (3.5 s per trial), but not during stimulus presentation (leaving a period of 1s per trial to record EEG data without tACS artifact, -600:100 ms relative to stimulus onset/offset). After each block, participants completed a questionnaire designed to quantify the intensity of perceived tACS side effects. Each trial started with the presentation of a central fixation dot for 1.2 s followed by a central Gabor patch for 0.3 s (Figure 1 B). Participants had to indicate its orientation as fast as possible by button press (Figure 1 C). After stimulus offset, the fixation dot reappeared and changed color as soon as participants gave their response (green for correct responses, red for incorrect responses; feedback duration 3 s). All three colors of the fixation dot were isoluminant and participants were instructed to maintain central fixation throughout the whole experiment. The Gabor patch spanned 5° visual angle and was presented against a white noise background (120 Hz rate) for masking afterimages. Orientations of the Gabor patch were horizontal (0°), left leaning (45° clockwise), vertical (90° clockwise) and right leaning (135° clockwise). The resulting four-alternatives forced-choice (4AFC) task has been shown to elevate precision and efficiency in estimating psychometric parameters when compared to a 2AFC task (Hou et al., 2015; Vancleef et al., 2018). Prior to the experimental blocks, participants trained orientation/button – correspondences until they reached 100% accuracy. In the experimental blocks, Gabor patches were presented in random order. Prior to each trial, the contrast value of the upcoming stimulus was adaptively chosen to optimize the estimation of parameters (see *Psychometric procedure*). Contrast varied between 0.05 and 0.5 Michelson contrast in 20 equidistant steps. A total number of 100 trials was presented in each tACS condition.

**Figure 1.**
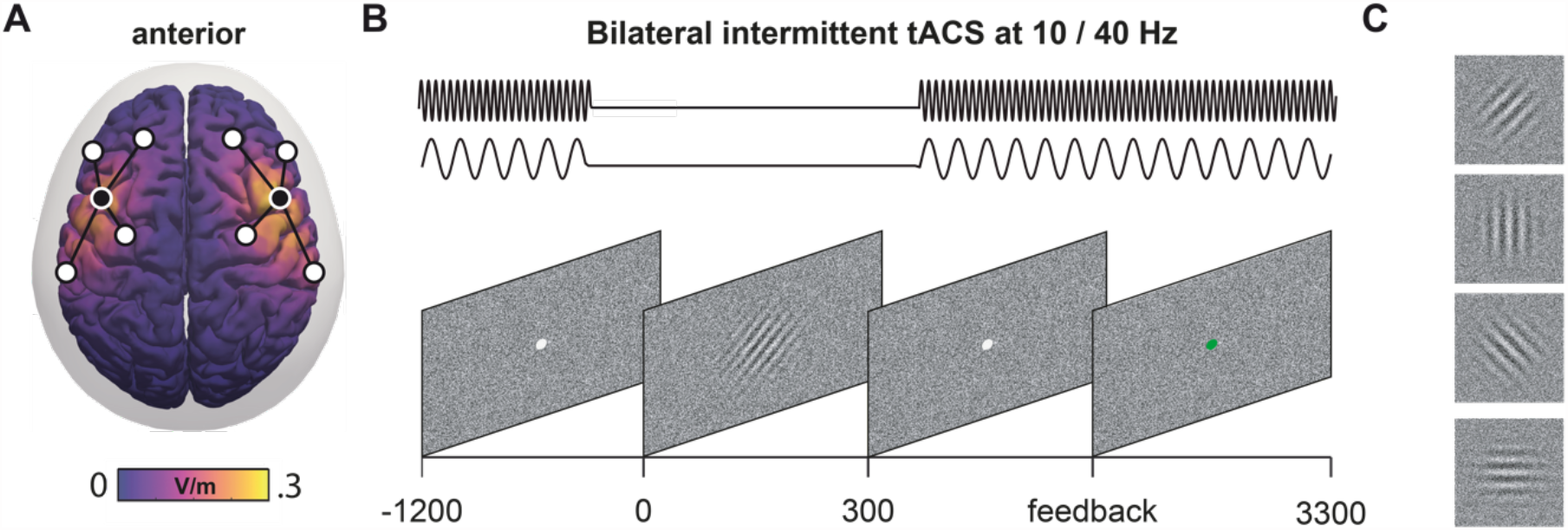
Experimental design. **(A)** Montages of tACS electrodes centred over left and right premotor cortices. Electrodes with different polarities are shown with inverted black/white color-coding. **(B)** Schematic of intermittent stimulation with respect to the trial progression. Alpha (10 Hz) or gamma (40 Hz) stimulation was omitted during the presentation of the stimulus presentation and the period directly before (−600:400 relative to stimulus onset). **(C)** Garbor patches with four different orientations were used as visual stimuli.

### Psychometric procedure

Psychometric functions were estimated using a Bayesian adaptive procedure (Prins, 2013). The psi-marginal adaptive method estimates slope and threshold of a psychometric function while allowing to adaptively target nuisance parameters lapse rate and guess rate if such estimation reduces uncertainty with respect to the parameters of primary interest. In short, a posterior distribution is defined by all free parameters and used for the selection of the next to-be-presented stimulus intensity. This selection is based on the expected information gain in the marginal posterior distribution defined across all parameters of interest. Prins (2013) found that estimating lapse rate as a nuisance parameter increased precision and reduced bias in estimating threshold and slope parameters. Accordingly, we fixed the guess rate at .25 and estimated primary parameters threshold and slope as well as the nuisance parameter lapse rate.

### Electric stimulation

Multi-electrode tACS was administered with two separate stimulators (DC-Stimulator Plus, Neuroconn, Germany) in external mode controlling one electrode montage each placed over the left or right premotor cortex. The external mode allows to control current output via voltage input. Voltage signals were generated in Matlab (5000 Hz sampling rate) and produced as an analog signal by a NI-DAQ device run with Labview (NI USB 6343, National Instruments, USA). For each montage, we used five Ag/AgCl ring electrodes (12 mm diameter) arranged in 4-in-1 multi-electrode montages (Patel et al., 2009). These montages have been shown to be optimal when aiming to restrict electric field spread (Saturnino et al., 2015). Here, we targeted bilateral premotor cortex (Vernet et al., 2014) and chose a montage with maximum field strength within this area based on simulations (see *Electric field simulation*, Figure 1 A). In order to get an evenly distributed electric field, we carefully prepared electrodes with equal impedances below 50 kΩ. In the beginning of each block, tACS was ramped up linearly from 0 to 2 mA (peak-to-peak) within 5 seconds. In sham blocks, stimulation was terminated thereafter (i.e., stimulators kept 0 mA output). In verum blocks, tACS continued with 2 mA (peak-to-peak) at a frequency of 10 or 40 Hz. Importantly, tACS was applied intermittently, meaning that tACS was omitted during stimulus presentation without ramping (−0.6 s and +0.1 s around stimulus onset).

### Electric field simulation

Electric fields were estimated using exact low-resolution electromagnetic tomography (eLORETA; Pascual-Marqui et al., 2011). We used a boundary-element three-shell head model (Nolte and Dassios, 2005) in combination with a cortical grid (MNI152). Electric field strength was estimated as the sum of linear combinations between eLORETA leadfield 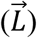 and injected currents at the stimulation electrodes (*α*_*i*_) for each cortical location 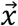:

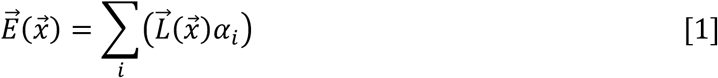

### Pupillometry

Pupil size of both eyes was tracked continuously with head-mounted glasses (Pupil Labs, Berlin, Germany) and recorded using PupilRecorder Software with default settings (Kassner et al., 2014). Pupil data was processed offline in Matlab (Mathworks, Massachusetts, USA) with custom scripts. For data analysis, data from one eye was chosen based on the cumulated confidence ratings of both eyes (Kassner et al., 2014) and pupil data was resampled at 20 Hz. Identification of blinks was based on the confidence rating (cutoff = .95) and pupil slope (cutoff = 5). For complete removal of blinks, identified blink epochs were padded on both sides with 200 ms and data was removed. Interpolation was performed with cubic spline interpolation on the whole time series. For the analysis of phasic pupil responses, data was filtered using a 3^rd^ order Butterworth filter (high-pass: .2 Hz, low-pass: 4 Hz). Finally, data was up-sampled with cubic spline interpolation to 1000 Hz, cut into epochs of single trials and z-transformed. The first trial was omitted. Further, we rejected trials containing at least one sample that exceeded three standard deviations of all sample points (mean 10±6 trials). Prior to analysis, pupil data was stratified to ensure that all conditions held the same number of trials in all conditions within participants.

## Statistics

### Mixed model analysis

Psychometric parameter estimates for threshold, slope and lapse rate were subjected to linear mixed model analysis using SPSS (IBM SPSS Statistics for Macintosh, Version 27.0.). We accounted for the influence of side effects and its interindividual variance by including measures of side effects, both subjective and objective, as covariates and by modeling random intercepts. A subjective account of side effects was captured by administering a questionnaire after each stimulation block that asked for skin sensations, pain and phosphenes (for details, see Misselhorn et al., 2019). As an objective measure of arousal that might be increased due to tACS, we measured pupil dilation (Kyle and McNeil, 2014; Murphy et al., 2014). In order to reduce the continuous pupil data to one value per trial, we averaged data from the period without tACS and consecutively averaged over trials. To avoid model overfitting, we opted for a minimum number of model parameters. The main fixed effect of interest was *stimulation* (sham, alpha, gamma). Covariates and interactions of covariates with *stimulation* were assessed beforehand and added only if relevant. That means, we run repeated measures ANOVAs with factor *stimulation* for each measure of side effect (skin, pain, phosphene and pupil) separately. Then, we added the side effect as a fixed main effect and interaction with *stimulation* to the mixed-model analysis only if the *p*-value of *stimulation* in the ANOVA was below 0.05. Please note that we used Friedmann’s ANOVA for rank data (skin, pain, phosphene) and parametric ANOVA for continuous data (pupil). Post-hoc comparisons were performed with paired *t*-tests or Wilcoxon signed rank tests. Subsequently, we reduced the mixed models by removing fixed effects that resulted in *p*-values above 0.05. Specifically, we removed non-significant effects starting with the lowest F-values. Main effects were only removed when they were not included in interactions. In order to quantify fit of the complete and reduced models, we computed Bayes information criterion (BIC). For the covariance structure, we assumed diagonal for fixed effects and identity for random effects (default settings SPSS). Post-hoc tests for fixed main effects and interactions were performed by pairwise multiple comparisons. To all post-hoc tests the significance level was set to α = .05 and we applied Sidak correction to control the family-wise error rate for multiple comparisons. In order to facilitate interpretation of the interactions, we performed bivariate Pearson’s or Spearman correlations depending on the scale of measurement.

### Permutation statistics

After model reduction, we used the resulting linear mixed models to compute marginal means for the assumption of no side effects. That is, covariates were set to zero (skin sensations or pain) or the average value (pupil). These estimations of parameters with reduced influence of side effects were used to reconstruct psychometric functions for each stimulation condition. Direct statistical comparison of the psychometric functions as a whole was performed by subtracting psychometric functions pairwise (alpha-sham, gamma-sham, alpha-gamma). Confidence intervals were constructed by means of non-parametric permutations statistics. That means, in 10,000 permutations, we randomly reassigned condition labels within participants and computed individual psychometric functions. Condition averages were subtracted and stored as part of the null distribution. The psychometric functions reconstructed from marginal means were then compared to a confidence interval spanning between the 0.83^th^ and 99.17^th^ percentile of the null distribution (two-tailed testing, α= 0.05, Bonferroni-corrected).

## Results

Prior to mixed model analysis of psychophysical parameters of contrast perception, we assessed the potential contribution of tACS side effects to perceptual modulations (Figure 2). In repeated measures ANOVA, we found subjective reports of skin sensations and pain to differ between stimulation conditions as well as pupil diameter, whereas reports of phosphenes did not show a significant association with tACS. Table 1 lists results from repeated-measures ANOVA as well as results from the post-hoc pairwise comparisons where applicable (values in the last three rows correspond to the average difference in ratings respectively pupil size between conditions, *p*-values were Bonferroni corrected). Skin sensations were elevated in both verum conditions compared to sham (alpha-sham: *p* = 0.003, gamma-sham: *p* = 0.012), while there was no difference between stimulation frequencies (alpha-gamma: *p* = 0.414). For pain, ratings were significantly increased only under gamma stimulation (gamma-sham: *p* = 0.009) but did not differ between sham and alpha stimulation (alpha-sham: *p* = 0.096) or between stimulation frequencies (alpha-gamma: *p* = 0.995). Pupil diameter prior to and during the presentation of the visual stimulus was largest during the gamma stimulation block and smallest for sham stimulation. Pairwise comparisons showed a significant increase from sham to gamma stimulation (gamma-sham: *p* = 0.009), but no differences between sham and alpha (*p* = 0.499) respectively alpha and gamma (p = 0.249). Following this analysis, we decided to include skin sensations, pain and pupil diameter as covariates in the mixed model of psychophysical parameters and modeled them as interactions with factor *stimulation* (sham/alpha/gamma).

**Table 1.**
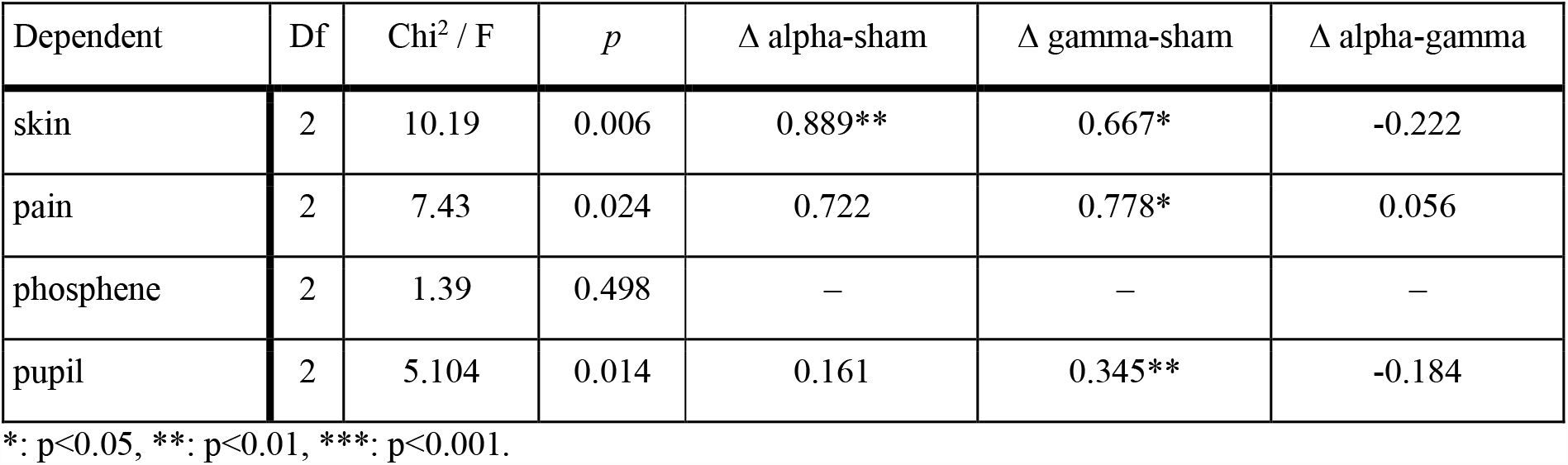
Repeated-measures ANOVA with factor stimulation (sham/alpha/gamma)

| Dependent | Df | Chi<sup>2</sup> / F | *p* | Δ alpha-sham | Δ gamma-sham | Δ alpha-gamma |
| --- | --- | --- | --- | --- | --- | --- |
| skin | 2 | 10.19 | 0.006 | 0.889** | 0.667* | -0.222 |
| pain | 2 | 7.43 | 0.024 | 0.722 | 0.778* | 0.056 |
| phosphene | 2 | 1.39 | 0.498 | — | — | — |
| pupil | 2 | 5.104 | 0.014 | 0.161 | 0.345** | -0.184 |

\*, $p < 0.05$ , \*\*, $p < 0.01$ , \*\*\*, $p < 0.001$ .

**Figure 2.**
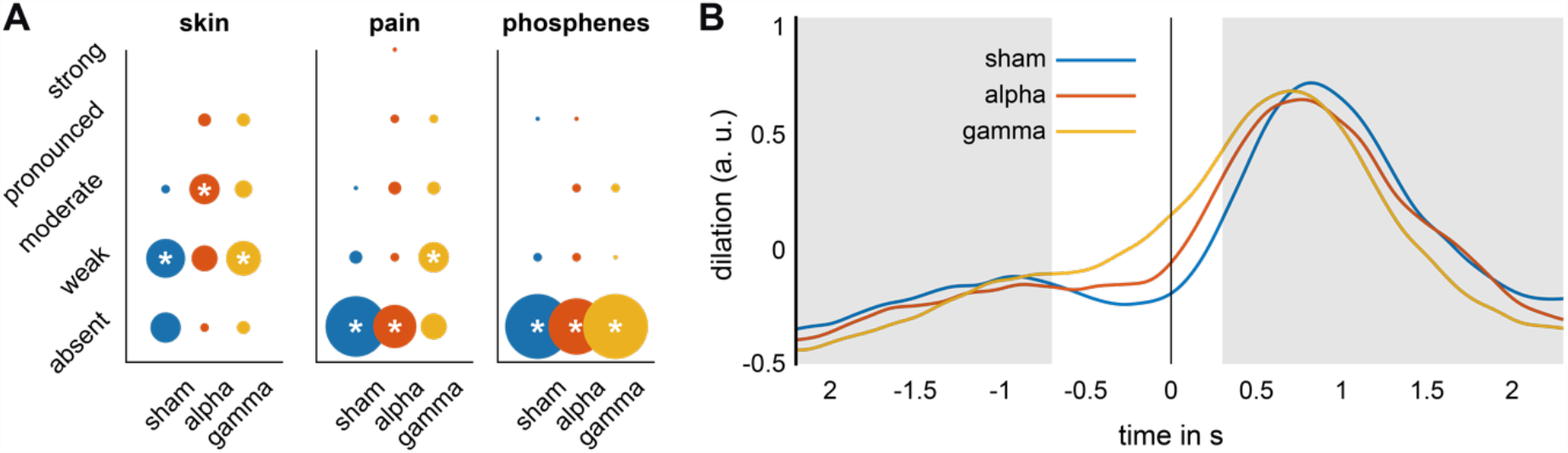
Side effects of tACS. **(A)** Ratings of perceived intensity of skin sensations, pain and phosphenes under sham (blue), alpha (10 Hz, red) and gamma stimulation (40 Hz, yellow). Circle size indicates the overall counts of ratings. Asterisks mark median responses. **(B)** Pupil dilation time-locked to the presentation of the stimulus under sham (blue), alpha (10 Hz, red) and gamma stimulation (40 Hz, yellow). Grey area indicates periods of tACS.

For the psychometric parameter threshold, mixed-model analysis revealed that threshold did not significantly differ between stimulation conditions (*p* = 0.934). Yet, it was positively associated with skin sensations, such that higher thresholds were measured in the context of stronger skin sensations (see Table 2). Post-hoc comparisons showed that this positive association was largely driven by sham and alpha stimulation, being significantly smaller during gamma stimulation (sham-gamma: *t(15*.*537)* = -3.799, *p* = 0.006; sham-alpha *t(23*.*599)* = - 2.075, *p* = 0.140; alpha-gamma: *t(16*.*229)* = 0.119, *p* = 0.999). Model fit improved from the basic model (only stimulation as fixed effect, no random effect, no covariates) to the final reduced model (BIC: -197.888/-200.690).

**Table 2.** Reduced linear mixed models for all three psychometric parameters.

| Source | dfn | dfd | *F* | *p* |
| --- | --- | --- | --- | --- |
| **Threshold** |  |  |  |  |
| Constant term | 1 | 37.433 | 1098.226 | 0.000 |
| Stimulation | 2 | 16.321 | 0.069 | 0.934 |
| Skin | 1 | 39.100 | 8.781 | 0.005 |
| Stimulation*Skin | 2 | 16.055 | 7.220 | 0.006 |
| **Slope** |  |  |  |  |
| Constant term | 1 | 43.194 | 4193.944 | 0.000 |
| Stimulation | 2 | 31.180 | 5.150 | 0.012 |
| Pain | 1 | 31.245 | 17.458 | 0.000 |
| Pupil | 1 | 42.435 | 10.611 | 0.002 |
| Stimulation*Pain | 2 | 31.852 | 3.882 | 0.031 |
| Stimulation*Pupil | 2 | 33.528 | 4.078 | 0.026 |
| **Lapse rate** |  |  |  |  |
| Constant term | 1 | 18.859 | 397.346 | 0.000 |
| Stimulation | 2 | 19.065 | 0.908 | 0.420 |

Slope was significantly influenced by stimulation condition (*p* = 0.012), with alpha stimulation significantly decreasing slope (alpha-sham: *t(28*.*264)* = -2,610, *p* = 0.041; alpha-gamma: *t(29*.*552)* = -3.050, *p* = 0.015). There was no significant difference between sham and gamma stimulation (*t(29*.*512)* = 0.652, *p* = 0.889). Additionally, pain and pupil size modulated slope in opposite direction. While reports of stronger pain were inversely related to slope, large pupil diameters were measured when slope was high. These effects depended on the stimulation condition. The influence of pain on the slope parameter was comparable between gamma and sham conditions (*t(22*.*559)* = 1.564, *p* = 0.346), but significantly lower in alpha compared to sham condition (*t(22*.*727)* = 2.656, *p* = 0.041). Alpha and gamma conditions did not show a significant difference with respect to the influence of pain on slope (*t(29*.*995)* = 1.675, *p* = 0.281). Pupil size showed a comparable positive association with slope induced by alpha and gamma stimulation (*t(27*.*435)* = -0.925, *p* = 0.742), but showed a negative association with slope under sham stimulation (gamma-sham: *t(28*.*604)* = 2.853, *p* = 0.024; alpha-sham: *t(29*.*754)* = 1.616, *p* = 0.312). Slope did not depend on the random variable subject which was therefore removed. Model fit of the reduced model improved slightly compared to the complete model (BIC: -57.922/-59.523). Finally, lapse rate did not depend on any fixed effect, but model fit slightly improved when modeling random intercepts (BIC: -280.058/-281.038).

Summarizing the main research question, we found that alpha stimulation significantly decreased psychometric slope compared to sham and gamma stimulation, while threshold and lapse rate parameters were unaffected. In order to detail contrast-dependent effects across the perceivable range, we planned to directly compare psychometric functions between stimulation conditions by subtraction and evaluate resulting difference functions with non-parametric permutation statistics (see Methods). However, as we demonstrated above, neurosensory side-effects influenced psychometric parameters. To control their influence in this analysis, we computed marginal means of the psychometric parameters with the mixed models by setting the questionnaire covariates equal to zero (i.e., absent skin sensation or pain) and the pupil covariate to the mean value. Descriptively, accounting for side-effects decreased thresholds and increased slopes in all three stimulation conditions. These parameter changes were of small magnitude and did not result in significant t-tests between corrected and uncorrected parameters (for all, *p* > 0.10; Figure 3 A). Yet, when comparing psychometric functions by more sensitive non-parametric permutations statistics, changes by side-effect correction were significant under both sham and gamma stimulation (Figure 3 B-D).

**Figure 3.**
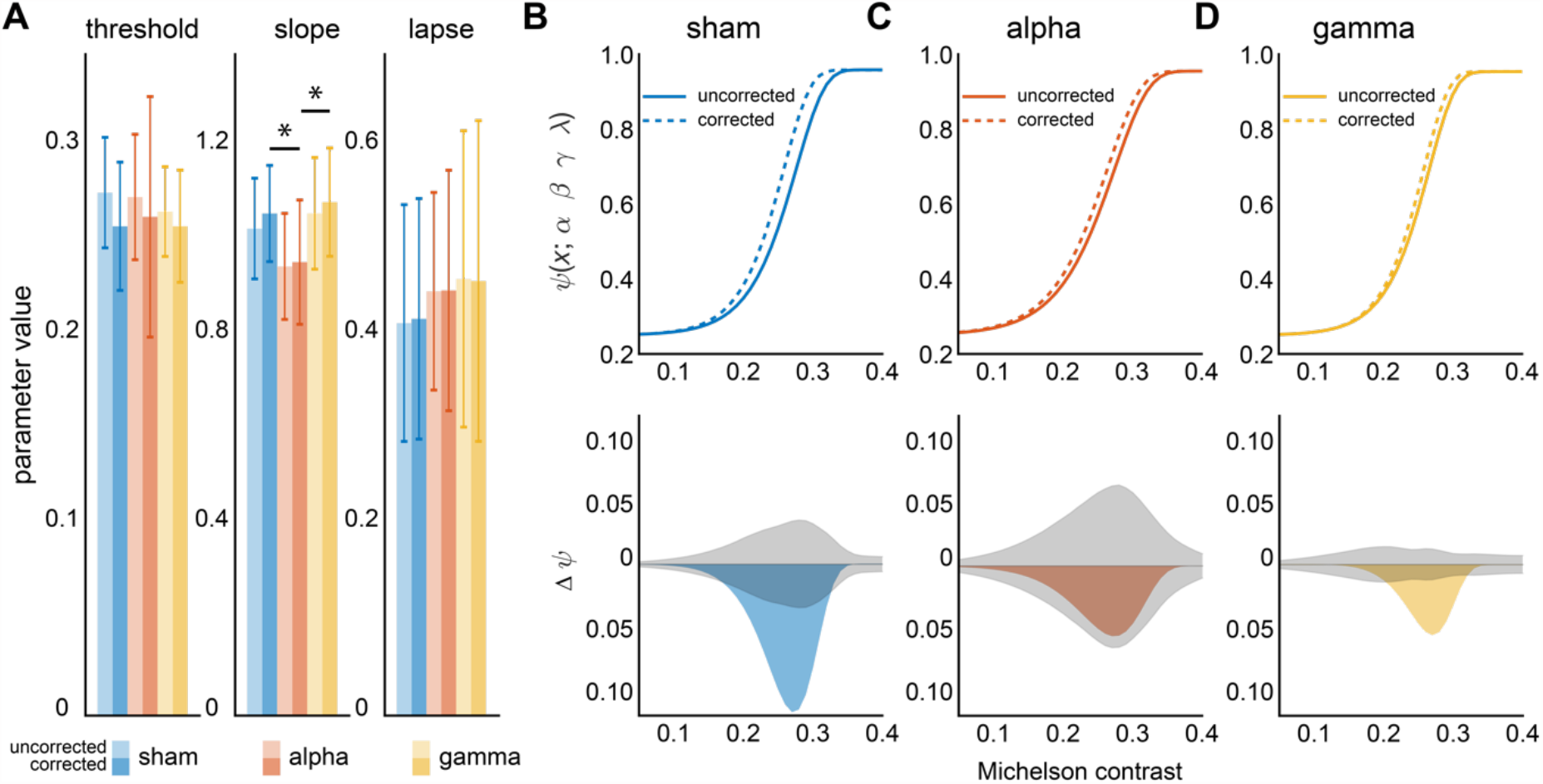
Side-effects of tACS on contrast perception. **(A)** Psychometric parameters of uncorrected (lighter colors) and corrected (darker colors) data. Slope was decreased by alpha stimulation before and after correction (asterisks). In the comparison between corrected and uncorrected data, we did find no significant differences. **(B-D) (top)** Psychometric functions of uncorrected and corrected data. **(bottom)** When focusing on psychometric functions as a whole by using non-parametric permutation statistics, data show significant differences between uncorrected and corrected data for sham and gamma stimulation. Gray area depicts 95% confidence interval as determined by non-parametric permutation statistics.

Finally, we applied the non-parametric statistics to the comparison between stimulation conditions to analyze contrast-dependent transcranial effects of stimulation. When reconstructing functions based on uncorrected estimations of psychometric parameters, we did not find significant differences between tACS conditions (Figure 4 A,B). Importantly however, when comparing corrected psychometric functions between stimulation conditions, we found a clear-cut effect of alpha stimulation compared to both control conditions, such that sub-threshold contrast perception was enhanced at the expanse of super-threshold perception (Figure 4 C,D). This pattern resulted from an isolated decrease in slope with unchanged threshold or lapse rate (Table 2). Please note that this pattern of results is consistent with the mixed model results of the whole data. Thus, correction for side effects did not alter the pattern of results overall, but accounts for inter-individual variability in stimulation outcome and, thereby, demonstrates the perceptual consequences of tACS effects with more detail and clarity.

**Figure 4.**
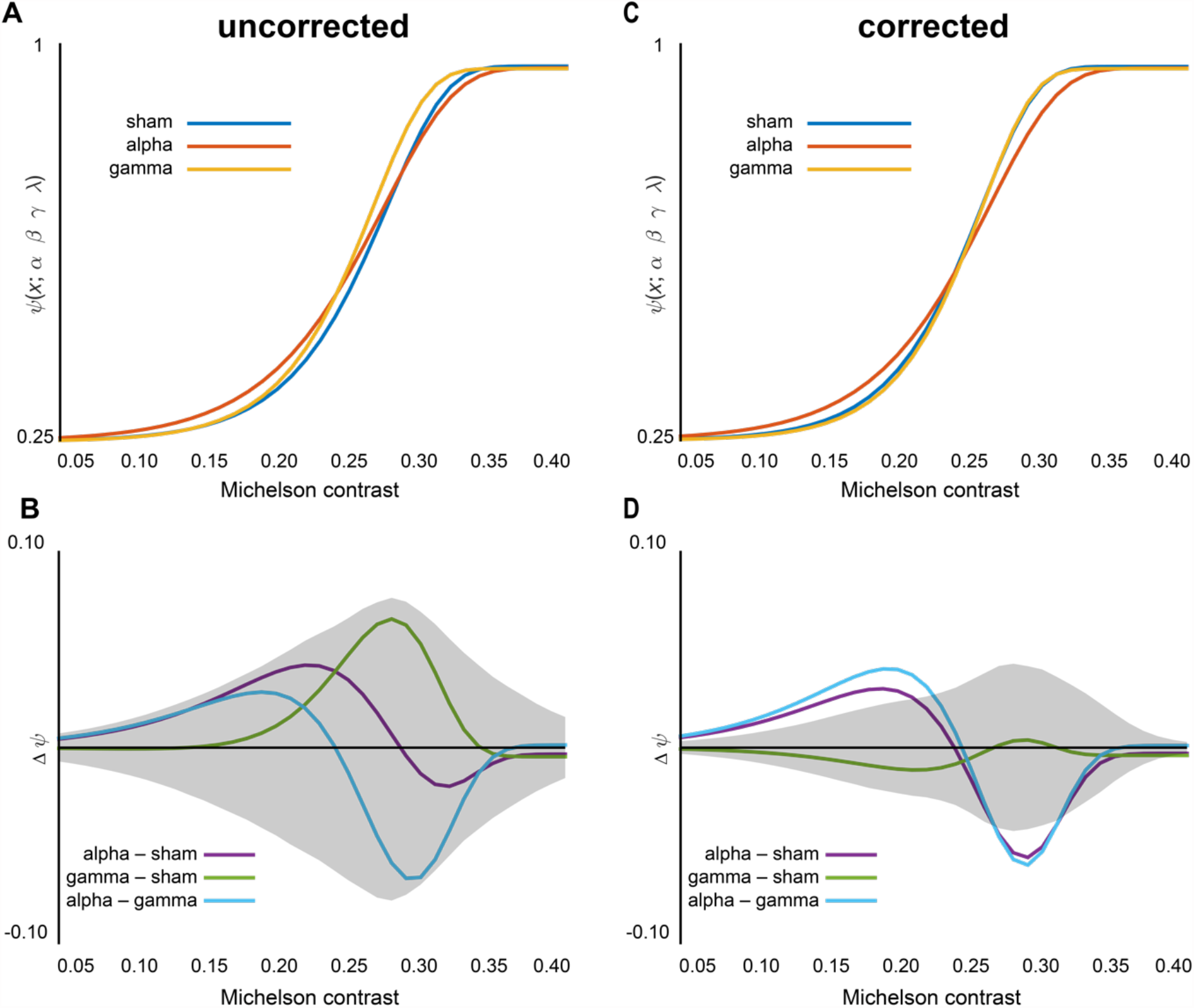
Effects of tACS on contrast perception. Upper row depicts psychometric function per stimulation condition and lower row depicts pairwise differences between psychometric functions and confidence intervals for these contrasts. **(A**,**C)** Uncorrected psychometric functions show trending effects for both alpha and gamma stimulation. Due to comparably high interindividual variability, none of the effects is statistically significant. **(B**,**D)** Corrected psychometric functions (reconstructed from marginal means of the mixed model) show a clear distinction between an effect for alpha, and no effect for gamma stimulation. The effect of alpha stimulation as well as the difference to gamma stimulation is statistically significant.

## Discussion

To investigate preparatory top-down influence on visual processing, we stimulated human premotor cortex with intermittent tACS at alpha (10 Hz) frequency during psychometric testing of visual contrast sensitivity and compared outcomes to gamma (40 Hz) and sham stimulation. Side effects of the electric stimulation were systematically measured and used in a linear mixed model analysis to control for potential transcutaneous and transretinal origins of perceptual effects. At the level of psychometric parameters of contrast perception, we find a significant effect of alpha tACS on the slope of psychometric functions. Yet, large variability of parameter estimates resulted in an unclear pattern when comparing psychometric functions directly by means of non-parametric permutation statistics, hampering the interpretation of stimulation effects. However, when correcting for neurosensory side effects of electrical stimulation, the effect of premotor alpha stimulation on the slope of the psychometric function presents with higher clarity: an improvement of low-contrast perception at the expanse of high-contrast perception.

Previous research on how attention can shape psychophysics of visual perception has shown both effects on threshold and slope (Cameron et al., 2002; Ling and Carrasco, 2006; Herrmann et al., 2010; Cutrone et al., 2014). While these studies consistently showed reduced thresholds due to attentional control, the effects on slope were mixed across studies and conditions. In some cases slope remained unchanged, resulting in a leftward shift of the psychometric function without changing its shape (Cameron et al., 2002; Herrmann et al., 2010). Other studies either showed an increase in slope (Ling and Carrasco, 2006) or a decrease (Cutrone et al., 2014), respectively. This inconsistency of findings might be related to different factors. First, task setting and related types of attention differed between studies. Collectively, these studies suggest that exogeneous attention might not involve a clear slope modulation, but that endogenous and feature-based attention do (Ling and Carrasco, 2006; Cutrone et al., 2014). Second, it was shown that spatial uncertainty and the relation between receptive field and stimulus size determine if the slope is modulated (Herrmann et al., 2010). Finally, it was suggested that response bias might also influence the perceptual effects of attention. In a series of experiments, Zhou and colleagues (Zhou et al., 2018) showed that attentional cueing in a same/different task selectively enhanced performance for low-contrast stimuli consistent with a combined effect of decreased slope and threshold. However, when properly controlling for the effects of response bias, the enhancement for low-level contrasts was accompanied by an attenuation for high level contrasts. This latter finding is consistent with our study where contrast-dependent improvement and attenuation of visual contrast perception resulted from an isolated change in slope. We suggest that the effect of frontal alpha tACS on slope mimics the induction of a functional state of endogenous attention in premotor cortex that improved feature-based selective attention.

Despite an abundance of work on psychophysics and its variability due to cognition, conclusive work on its neurophysiological underpinnings is missing. In work on perceptual learning in rhesus monkeys, it was demonstrated that thresholds as well as slopes decreased as monkeys learned a motion discrimination task (Gold et al., 2010). The authors showed that the changes in slope could be explained by an increasingly selective readout of sensory information due to (non-linear) pooling of sensory responses onto decision-related areas. They suggested that this process might be equally important in feature-based attention (Maunsell and Treue, 2006), and that the pooling might rely on a refinement of functional connectivity between sensory and decision-related areas (Gold et al., 2010). In humans, there is good evidence that improving interactions in cortical networks involves oscillations in the alpha/beta band that coordinate sensory transmission in the gamma band (Donner and Siegel, 2011; Bonnefond et al., 2017). The regulation of alpha oscillations has been investigated extensively in the context of spatial attention, where anticipatory attention was reliably related to a lateralization of alpha power in occipitoparietal cortex (e.g. Jensen and Mazaheri, 2010). Recent studies showed that the degree of lateralization depended on activity in premotor cortex and the strength of frontoparietal structural connectivity via the superior longitudinal fasciculus (Marshall et al., 2015a), suggesting that premotor cortex is driving the up- and down-regulation of alpha oscillations in occipitoparietal cortex. This idea was confirmed in a TMS study where stimulation of the premotor cortex modulated anticipatory alpha lateralization in occipitoparietal cortex in a spatial attention task (Marshall et al., 2015b). In another experiment, it was demonstrated that such regulation of occipitoparietal alpha activity could be due to oscillatory alignment between occipital and frontal alpha oscillations (Veniero et al., 2021). It is, thus, possible that improved selective readout from sensory neurons due to attentional control is implemented by modulating functional connectivity between frontal and occipital cortex in the alpha band. In fact, it was shown that functional networks instantiated via nested alpha and gamma oscillations establish communication serving top-down control of sensory gating (Bonnefond and Jensen, 2015; Bonnefond et al., 2017). We suggest that tACS over frontal cortex in our experiment increased neuronal activity in the alpha band, which was shown for different brain regions in ferrets (Huang et al., 2021), monkeys (Krause et al., 2019; Johnson et al., 2020), and humans (Schwab et al., 2019; Fiene et al., 2020). Secondary to these local changes in oscillatory synchronization and/or amplitude, it is reasonable to assume changes in functional connectivity to connected brain regions (Wischnewski et al., 2023). Only recently, such an indirect effect of electric stimulation has been suggested to explain event-related potential modulations in premotor cortex that were induced by alpha tACS targeting the parietal cortex during a visuo-spatial attention paradigm (Radecke et al., 2023). Taken together, we speculate that in this study tACS helped to instantiate functionally relevant fronto-occipital alpha networks underlying selective attention and, thereby, enhanced a non-linear process of sensory readout from visual areas.

In addition to the effects of tACS as the primary outcome, we investigated side effect characteristics of tACS and how they influence contrast perception. Although we only found weak-to-moderate (skin sensations) or absent-to-weak (pain and phosphenes) group level reports of tACS-related side effects, we found a significant impact of these side effects on parameter estimates of psychometric functions in the mixed models. Psychometric threshold of contrast perception was increased by strong skin sensations which could be due to a cross-modal shift of attention away from the task-relevant visual modality towards the salient somatosensory stimulus (Misselhorn et al., 2016). Interestingly, not only verum but also sham stimulation led to an increased threshold if skin sensations were subjectively perceived as moderate-to-strong. This finding suggests that, independent of the actual flow of current through skin and peripheral nerves, participants expected stimulation and sometimes imagined corresponding side effects or perceived actual tactile sensation (e.g., itching) that was unrelated to the electrical stimulation. Imagined or perceived, both types of subjective side-effects can induce shifts of attention at the expanse of contrast perception. This idea is further illustrated by the in-depth analysis of psychometric changes after mixed model correction. When comparing psychometric functions as a whole using non-parametric statistics, we find that mixed modeling reduced the impact of side-effects. Note, however, that this impact on parameter estimates was subtle, such that parametric statistics did not detect significant changes between uncorrected and corrected parameters. Next to threshold, also slope was associated with side effects on pain perception and pupil diameter. Pupil diameter was generally increased under verum stimulation possibly indicating an overall increase in arousal that has been shown to affect subcortical and cortical processing (Joshi et al., 2016; Sharon et al., 2021; Pfeffer et al., 2022). More specifically, pupil-linked arousal was shown to be associated with noradrenergic neuromodulation by the locus coeruleus that mainly affected activity in parietal and pre-frontal cortices, but did not boost sensory responses (de Gee et al., 2017). In accordance with this idea, and in support of our previous claim, effects on the slope parameter might rather reflect top-down influence on visual processing and possibly decision-making processes.

Taken together, our data suggest an important role for premotor alpha oscillations in the control of endogenous attention. We propose that premotor alpha tACS mimicked a state of endogenous attention and helped instantiating a functional fronto-occipital network in the alpha band. We speculate that this network modulation allowed for improved sensory readout from visual cortex which resulted in a decrease of psychometric slope, effectively broadening the dynamic range for contrast perception. Additionally, we demonstrated that side-effects of during transcranial electric stimulation studies have complex effects on perception which, however, can be controlled for with linear mixed modeling. This approach can be of use to the field of non-invasive brain stimulation in general by providing a powerful tool to handle inter-individual variability in stimulation outcome.

## Acknowledgments

This work was funded by the German Research Foundation (DFG) (SFB 936-178316478-A3 awarded to A.K.E. and T.R.S.; SPP 1665/EN 533/13-1 awarded to A.K.E; SPP 1665/SCHN 1511/1-2 awarded to T.R.S; JOR was funded by LE 1122/7-1), by the EU (ERC-2022-AdG-101097402 awarded to A.K.E.) and by the Studienstiftung des deutschen Volkes (awarded to M.F.).

